# Glucose-dependent signalling pathways regulate TE differentiation in bovine embryos

**DOI:** 10.64898/2026.04.08.717294

**Authors:** Jinlong Qiu, Roger Sturmey

## Abstract

Glucose uptake and metabolism increase markedly during the transition from morula to blastocyst. However, the functional roles of glucose during this process in bovine embryos remain incompletely understood. Here, we demonstrate that glucose specifically modulates trophectoderm (TE) differentiation without affecting the inner cell mass (ICM), acting through Hippo signalling via the hexosamine biosynthetic pathway (HBP) and the pentose phosphate pathway (PPP) rather than glycolysis.

Consistently, scRNA-seq analysis reveals that unlike glycolysis-related genes, which show only stage-specific changes, HBP- and PPP-related genes exhibit significant differential expression between the TE and ICM lineages. Furthermore, we found that glucose deprivation does not impair pyruvate uptake or intracellular pyruvate levels in blastocysts, yet it significantly depletes the terminal metabolites of the HBP and PPP. Together, these findings reveal that glucose-dependent regulation of TE differentiation is conserved in bovine embryos.

## Introduction

Glucose is a key metabolic substrate during early embryonic development [1, 2]. Classical models of glucose metabolism presumed that glucose is predominantly catabolized through glycolysis to generate pyruvate, which enters the tricarboxylic acid (TCA) cycle to satisfy the energetic requirements of cell proliferation and development [3, 4].

However, recent studies in mouse embryos have challenged this traditional bioenergetic view of glucose metabolism [5-7]. These studies demonstrate that, during early preimplantation development, glucose contributes only minimally to glycolytic flux and ATP production [5-7]. Instead, glucose-derived carbon is preferentially channeled into HBP and PPP, supporting biosynthetic processes and redox homeostasis [5, 7].

In particular, HBP-derived O-linked N-acetylglucosaminylation (O-GlcNAc) has been implicated in the regulation of the Hippo signaling pathway, thereby influencing TE-associated transcriptional programs [5, 8, 9]. In parallel, the PPP has been shown to function upstream of mechanistic target of rapamycin (mTOR) signaling to regulate TE specification [5, 10, 11]. Whether this glucose-driven signaling paradigm is conserved across bovine embryogenesis, however, remains to be elucidated.

In this study, we demonstrate that while glucose-mediated regulation of the TE program is conserved in bovine embryos, their metabolic dependency is significantly more intricate than that observed in murine models. Notably, bovine embryos exhibit a heightened tolerance to glucose deprivation, retaining the capacity for blastocyst formation even in its absence. Furthermore, the contributions of the HBP and PPP appear highly context-dependent, with developmental outcomes varying according to the timing of inhibition and the specific metabolic nodes targeted. Collectively, our findings underscore a critical role for glucose in the TE formation of large mammals while revealing the nuanced metabolic architecture underlying this process.

## Materials and methods

Unless specified otherwise, all reagents and chemicals were obtained from Merck (Dorset, UK).

### In vitro bovine embryo production

In vitro maturation (IVM), fertilization (IVF), and culture (IVC) of bovine embryos were performed according to previously described protocols with minor modifications [12-14]. IVM, IVF, and SemenPrep media were obtained from IVF Bioscience (Falmouth, UK). Bovine ovaries were harvested from animals slaughtered for commercial purposes unrelated to the study. Ovaries were collected and transported to the laboratory within 2 hours of slaughter. Cumulus–oocyte complexes (COCs) were collected from 3–8 mm follicles on the ovarian surface. COCs were cultured in IVM medium at 39°C, 5% CO_2_ for 22 hours. Following maturation, groups of 25–30 COCs were co-incubated with spermatozoa (Genus Plc) (1–5 × 10^6^/mL) purified from frozen–thawed semen using SemenPrep medium. IVF was carried out at 39°C, 5% CO_2_ for 10–14 hours. After fertilization, cumulus cells were removed. Presumptive zygotes were subsequently cultured in BO-SOF medium supplemented with either 0 mM or 1.5 mM glucose at 39 °C, 5% CO_2_ and 5% O_2_.

The BO-SOF formulation was prepared according to the protocol described in our previous publication [15, 16]. The inhibitors used in this study are listed in Supplementary Table 1. Unless otherwise stated, inhibitors were dissolved in BO-SOF and added from E0. The inhibitor concentrations were selected based on previously published studies [5, 17].

### Metabolic assay

Morula-stage embryos were individually cultured for 24 hours in 5-μL microdrops, after which embryos that successfully developed to the blastocyst stage were identified and their spent culture media were collected. All samples were immediately stored at −78°C until subsequent metabolic assays. Pyruvate concentrations were quantified using enzyme-coupled fluorometric methods adapted from a previously established protocol [15]. All enzymes used for metabolic analyses were obtained from Roche (Burgess Hill, UK). Standard curves were generated from serial dilutions of freshly prepared metabolite standards. Fluorescence values were calculated as the difference between pre- and post-incubation readings and converted to absolute concentrations. All measurements were performed in technical triplicate, and assays with standard curve R^2^ values < 0.98 were excluded from further analysis.

### Measurement of Basal Oxygen Consumption Rate (OCR)

The OCR detection protocol is based on previously published articles [16]. Briefly, OCR was assessed using a Seahorse XFp Analyzer (Agilent Technologies). Seahorse analysis was performed following the manufacturer’s recommendations for OCR. Embryos were allocated in small groups (6-10 bovine blastocysts per well) in 180 µl of HEPES-buffered SOF medium appropriate for the developmental stage.

### JC-1 staining

JC-1 (Thermo Fisher) was initially dissolved in DMSO and subsequently diluted to a final concentration of 1 µM in SOF medium. Embryos were incubated in this solution for 30 minutes under standard culture conditions. Following three washes with HEPES-buffered SOF, the embryos were imaged using a Zeiss LSM710 confocal microscope.

### Immunofluorescence (IF)

Samples were first washed three times with 0.1% polyvinylpyrrolidone/phosphate-buffered saline (PBS), then fixed with 4% paraformaldehyde/PBS for 30 minutes and then permeabilized with 0.5% Triton X-100/PBS for 30 minutes at room temperature. The blocking step was performed in blocking buffer (PBS containing 10% FBS and 0.1% Triton X-100) for 1 hour. Afterwards, samples were incubated overnight with primary antibodies in blocking buffer at 4°C. After three washes, embryos were incubated with secondary antibodies for 2 hours. Finally, samples were stained with Hoechst3342 (Hoechst) (Merck) for 15 minutes and then mounted. Images were acquired using inverted fluorescence microscope (Zeiss AXIO Vert.A1) or a laser scanning confocal microscope (Zeiss LSM 710) with a 40x objective lens.

FIJI (https://imagej.net/software/fiji/) was used for image visualization, cell counting, and signal intensity measurements. Depending on the experiment, signal intensity was measured and background subtracted to analyze absolute intensity. All antibodies (Supplementary Table S2 and S3) were validated to react with bovine cells [12, 18, 19].

### Single-cell RNA-seq data acquisition and processing

The single-cell RNA sequencing dataset of bovine preimplantation embryos analyzed in this study was obtained from the publicly available Gene Expression Omnibus (GEO) database under accession number GSE239782 [20]. This dataset contains transcriptomic profiles of bovine embryos at defined developmental stages, including morula, early blastocyst, mid-blastocyst, and late blastocyst stages. Raw count matrices were downloaded and processed using the Seurat R package (version 4.5.2). The calculation of module scores and all analysis codes were adapted from previously published studies [21].

### Statistical Analysis

Unless otherwise noted, all experiments were performed with at least three biological replicates. Differences between groups were evaluated using two-tailed unpaired Student’s t-tests. Graphs were generated with GraphPad Prism 10.

## Results

### Glucose plays a role in blastocyst formation

To investigate the functional requirement for glucose during bovine preimplantation development, embryos were cultured from embryonic day 0 (E0) in either glucose-free medium (G−) or control medium (CTRL). Subsequent monitoring revealed no significant differences in developmental rates between the G− and CTRL groups at the 4-cell (E2.0), 8–16-cell (E3.5), or morula (E5.0) stages (Fig. 1A, B). However, by E6.5, blastocyst formation rates were markedly reduced in the G− group (Fig. 1A, B). Furthermore, those few embryos that successfully reached the blastocyst stage failed to hatch normally by E8.5 (Fig. 1C, D). This morphological impairment was accompanied by a significant reduction in total cell count, a deficit that was already detectable as early as the morula stage (Fig. S1A–D). Collectively, these data suggest that while glucose is not strictly required for the blastocyst transition, it is indispensable for achieving optimal developmental efficiency, cellular proliferation, and subsequent hatching. Inhibition of glucose metabolism using the hexokinase inhibitor 2-deoxy-D-glucose (2-DG) similarly reduced blastocyst rate, confirming the important role of glucose in blastocyst formation (Fig. S1F, G).

**Fig 1.**
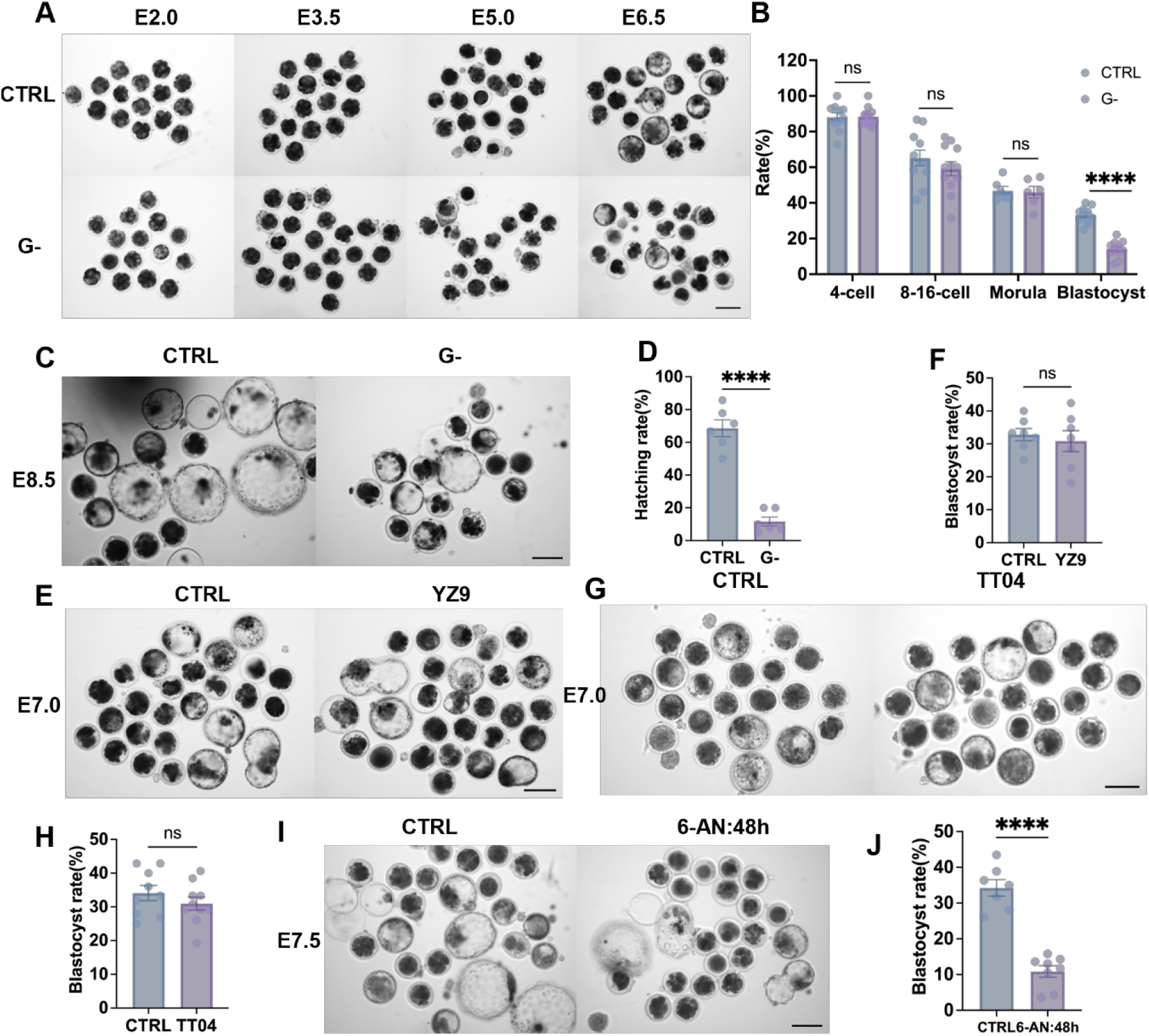
Glucose deprivation and inhibition of glucose metabolism impair bovine blastocyst formation. (A–D) Representative brightfield images of bovine embryos cultured in G− or CTRL media. Quantification of the proportion of embryos developing to the 4-cell (E2.0), 8–16-cell (E3.5), morula (E5.0) and blastocyst (E6.5) stages, and the proportion of hatching embryos at E8.5. (E, F) Representative images and corresponding blastocyst formation rates of E7.0 embryos cultured in the presence of the glycolysis inhibitor YZ9, relative to CTRL. (G, H) Representative images and blastocyst formation rates of E7.0 embryos cultured with the HBP inhibitor TT04, relative to CTRL. (I, J) Representative images and blastocyst formation rates of E7.5 embryos cultured with the PPP inhibitor 6-AN for 48 hours, relative to CTRL. All percentages were calculated relative to the total number of zygotes cultured per treatment. Data are presented as mean ± SEM from at least six independent replicates (≥15 embryos per group). Statistical significance was determined using a two-tailed unpaired Student’s t-test: ns, not significant; **P < 0.01; ****P < 0.0001. Scale bars, 50 μm.

To dissect the contributions of different glucose metabolic pathways, specific inhibitors were used to block glycolysis, HBP, and PPP (Fig. S1E). Inhibiting glucose flux into pyruvate with YZ9 had no effect on blastocyst formation rate (Fig. 1E, F). In contrast, blocking the PPP with 6-aminonicotinamide (6-AN) or the HBP with 6-Diazo-5-oxo-L-norleucine (DON) induced morula-stage developmental arrest (Fig. S1H, J), which could not be rescued by supplementation with their downstream metabolites (Fig. S1I, K). However, inhibition of the downstream HBP enzyme O-GlcNAc transferase (OGT) using ST045849 (TT04) had no effect on blastocyst formation rate (Fig. 1G, H). In addition, when PPP inhibition was delayed until E5.5, embryos were able to reach the blastocyst stage, although the blastocyst formation rate was significantly reduced (Fig. 1I, J). Together, these results demonstrate that glucose supports blastocyst formation through divergent metabolic nodes rather than through a uniform glycolytic requirement.

### Glucose metabolism regulates TE through Hippo pathway

To evaluate the impact of glucose on lineage specification, we performed immunofluorescence (IF) analysis on E7.0 blastocysts (Fig. 2A). Quantitative analysis showed that glucose deprivation significantly reduced CDX2 fluorescence intensity and the overall proportion of CDX2^+^cells within the blastocyst (Fig. 2B, C). Consistently, the expression of an additional TE marker, GATA3, was also diminished in the absence of glucose (Fig. S2A, B). In contrast, neither the protein levels of the ICM marker SOX2 nor the proportion of SOX2^+^cells were significantly altered (Fig. 2B, C). Similarly, the expression and cell count for the epiblast (EPI) marker NANOG and the primitive endoderm (PrE) marker GATA6 remained comparable between groups (Fig. 2D–F). These findings collectively indicate that glucose deprivation selectively impairs TE differentiation while leaving ICM lineage specification largely unaffected.

**Fig 2.**
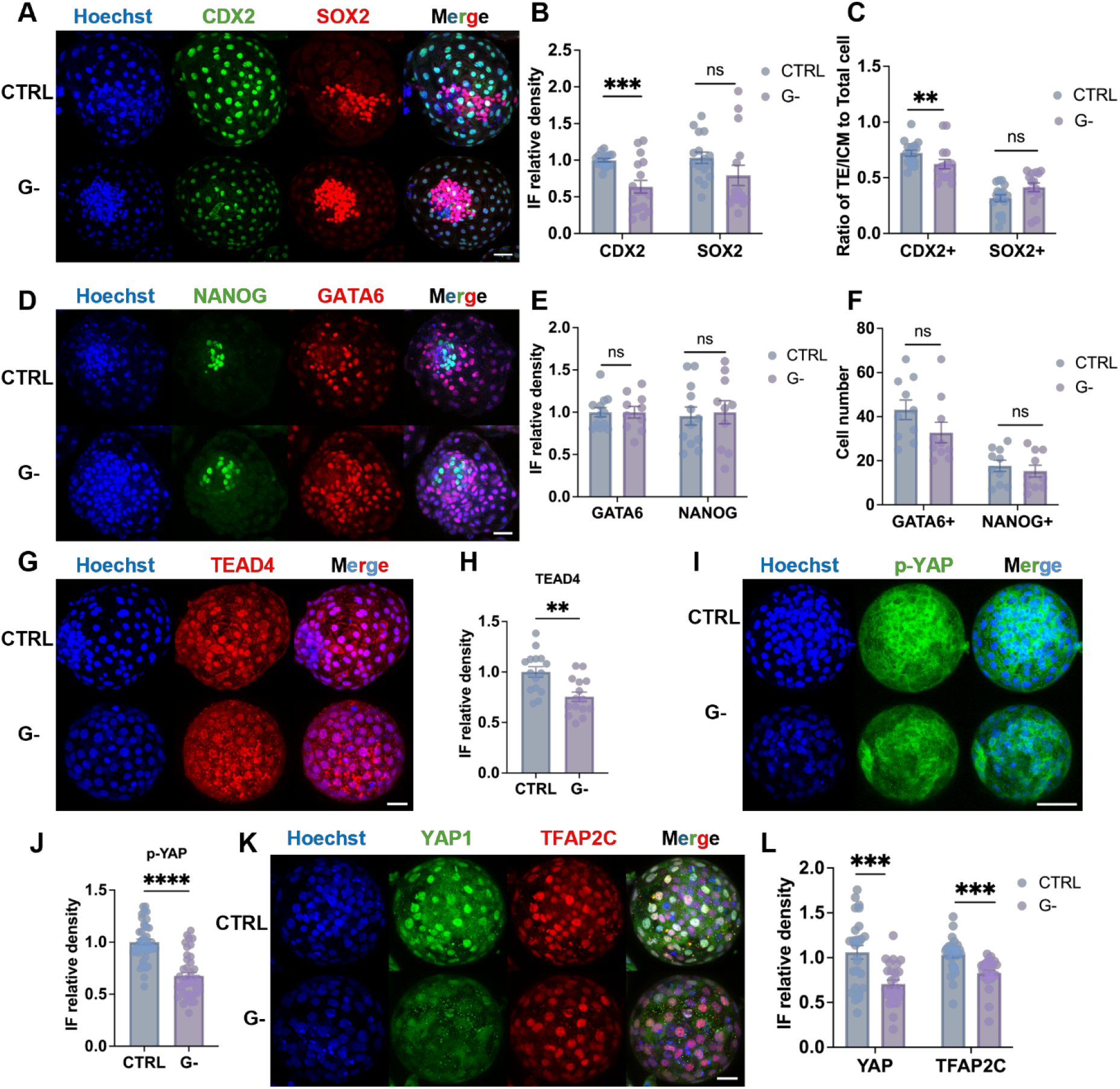
Glucose deprivation attenuates the Hippo signaling to impair TE specification. (A–C) Immunofluorescence staining and quantitative analysis of the TE marker CDX2 and ICM marker SOX2 in E7.0 bovine blastocysts cultured under CTRL and G− conditions. (D–F) Immunofluorescence staining and quantification of EPI marker NANOG and PrE marker GATA6 in E8.5 blastocysts cultured under CTRL and G− conditions. (G–L) Immunofluorescence staining and quantitative analysis of Hippo pathway components, including TEAD4 (G, H), p-YAP (I, J), YAP (K, L), and TFAP2C (K, L), in E7.0 blastocysts cultured under CTRL and G− conditions. For quantification, relative lineage-specific cell ratios were calculated for TE (CDX2^+^/Hoechst^+^), ICM (SOX2^+^/Hoechst^+^), EPI (NANOG^+^/Hoechst^+^), and PrE (GATA6^+^/Hoechst^+^) populations (n ≥ 3 biological replicates, with ≥3 blastocysts per group). Data are presented as mean ± SEM. Statistical significance was determined using a two-tailed unpaired Student’s t-test: ns, not significant; **P < 0.01; ***P < 0.001; ****P < 0.0001. Scale bars, 100 μm.

Given the central role of Hippo signalling in TE specification [22, 23], we next examined its activity in response to glucose availability. IF analysis revealed a significant reduction in phosphorylated YAP (p-YAP) levels in G− embryos (Fig. 2I, J). Consistently, the nuclear localization of YAP, as well as the abundance of TE-associated transcription factors TFAP2C and TEAD4, was markedly decreased under G− conditions (Fig. 2G, H, K, L). These results indicate that glucose levels modulate TE program execution via the Hippo-YAP axis. Similar phenotypes were observed following the pharmacological inhibition of glucose metabolism with 2-DG. Treatment with 2-DG significantly reduced total cell counts and shifted the TE-to-ICM ratio in E7.0 blastocysts (Fig. S2E), characterized by a selective suppression of CDX2 while leaving SOX2 levels unaffected (Fig. S2C–F). Furthermore, the expression of YAP, TFAP2C, and p-YAP was significantly diminished following 2-DG treatment (Fig. S2G–J), reinforcing the conclusion that glucose status specifically governs TE differentiation through the regulation of Hippo signaling components.

### Glucose availability regulates HBP activity and epigenetic landscapes

To identify the metabolic drivers of glucose-dependent TE specification, we performed a secondary analysis of publicly available metabolomics data from bovine blastocysts subjected to glucose-free culture [24]. While glucose deprivation led to a broad reduction in intermediates across several metabolic branches, the steady-state levels of pyruvate and ribose were unaffected, indicating that the outputs of glycolysis and the PPP are relatively preserved (Fig. S3A). In contrast, HBP metabolites, including N-acetylglucosamine (GlcNAc) and UDP-GlcNAc, were markedly reduced (Fig. S3A), highlighting a selective vulnerability of the HBP to glucose deprivation.

To validate our metabolomics findings, we first assessed mitochondrial membrane potential (ΔΨm) to assess the impact of glucose availability on cellular energy homeostasis. Glucose deprivation significantly reduced the orange/green JC-1 fluorescence ratio in bovine blastocysts, indicative of mitochondrial depolarization (Fig 3A, B). Interestingly, basal oxygen consumption rates (OCR), measured via Seahorse analysis, remained comparable between G− and CTRL embryos (Fig. 3C). This suggests that while glucose is essential for maintaining ΔΨm, bovine embryos possess the metabolic flexibility to sustain basal respiratory flux through the utilization of alternative substrates [25, 26].

**Fig 3.**
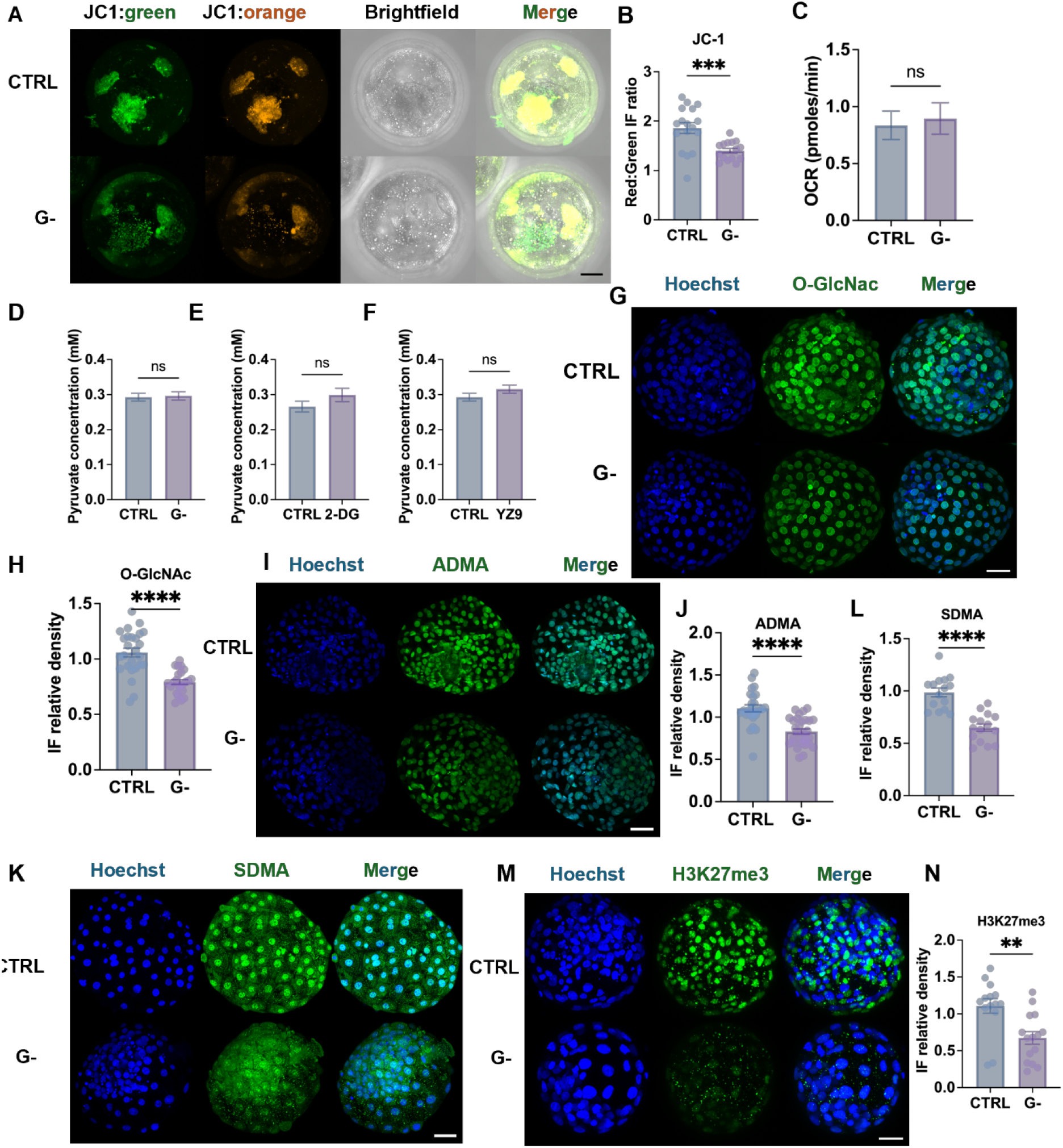
Glucose deprivation compromises mitochondrial function and epigenetic landscapes. (A, B) Representative fluorescence images of JC-1 staining in CTRL and G− blastocysts. Orange fluorescence indicates J-aggregates, whereas green fluorescence indicates J-monomers. Quantification of the red/green JC-1 fluorescence ratio. (C) OCR measured by Seahorse XF analysis in CTRL and G− blastocysts. (D–F) Pyruvate concentrations in culture medium following 24 h embryo culture under G−, 2-DG-treated, and YZ9-treated conditions, relative to CTRL. (G, H) Representative immunofluorescence images and corresponding quantification of global O-GlcNAc (green) levels in CTRL and G− blastocysts. (I-N) Representative immunofluorescence images and quantification of ADMA, SDMA, H3K27me3 in CTRL and G− blastocysts. Data are presented as mean ± SEM. Statistical significance was determined using a two-tailed unpaired Student’s t-test (n ≥ 3 biological replicates, with ≥5 blastocysts per group): ns, not significant; **P < 0.01; ***P < 0.001; ****P < 0.0001. Scale bars, 100 μm.

To determine whether embryos compensate for the absence of glucose by increasing reliance on exogenous pyruvate, pyruvate consumption from the culture medium was assessed. Measurement of pyruvate concentrations in spent culture media revealed no significant differences between CTRL and G-embryos (Fig 3D). Consistently, 2-DG or YZ9 inhibition did not significantly alter pyruvate concentration as well (Fig 3E, F). Collectively, these findings indicate that embryos do not trigger a compensatory increase in exogenous pyruvate uptake to offset the energy deficit caused by glucose deprivation.

To validate the impact of glucose withdrawal on the HBP identified in our metabolomics re-analysis, we assessed global O-GlcNAc levels. Consistent with the depletion of HBP intermediates observed in our metabolomics analysis, O-GlcNAc levels were significantly diminished in blastocysts cultured under G− conditions or treated with 2-DG (Fig. 3G, H and Fig. S3C, D). These results verify that restricted glucose availability directly compromises HBP-mediated protein O-GlcNAcylation in the developing embryo.

In parallel, our metabolomics re-analysis highlighted significant alterations in metabolites associated with amino acid methylation, specifically those linked to lysine trimethylation (Kme3) and arginine methylation (ADMA and SDMA) (Fig. S3B). Guided by these observations, we examined the relationship between glucose flux and epigenetic remodeling. Both glucose deprivation and 2-DG treatment markedly reduced ADMA and SDMA levels, which was accompanied by a pronounced decrease in the repressive histone mark H3K27me3 (Fig. 3I–N and Fig. S3C–J). These findings establish glucose availability as a key determinant of the epigenetic landscape in bovine blastocysts, likely through the regulation of methyl-donor availability [27].

### HBP-Dependent O-GlcNAc Regulates CDX2

To determine whether the HBP signaling axis operates through O-GlcNAc during bovine blastocyst development, we first verified that pharmacological inhibition of OGT (using TT04) significantly reduced global O-GlcNAc levels (Fig. 4G, H). We then assessed the impact of O-GlcNAc inhibition on lineage specification. TT04 treatment specifically downregulated CDX2 expression without perturbing SOX2 levels (Fig. 4A, B, while total cell number and the relative proportions of TE and ICM cells remained unchanged (Fig. 4C, D).

**Fig 4.**
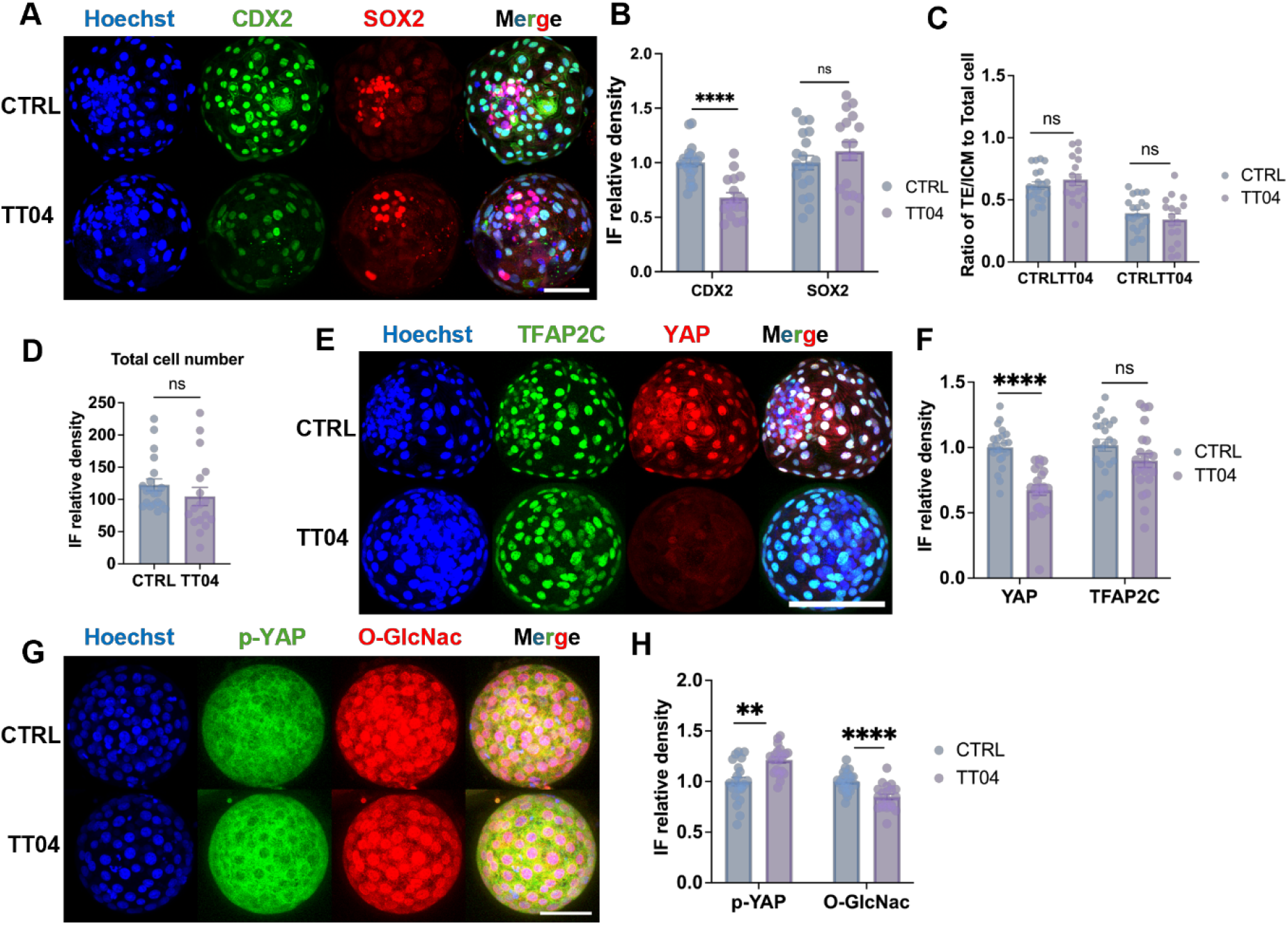
Inhibition of O-GlcNAc disrupts YAP and TE specification. (A, B) Representative immunofluorescence images and quantitative analysis of CDX2 and SOX2 in E7.0 embryos cultured under CTRL and TT04 treated conditions. (C, D) Quantitative analysis of total cell number (Hoechst^+^), and the proportions of TE (CDX2^+^/Hoechst^+^) and ICM (SOX2^+^/Hoechst^+^) cells in E7.0 embryos cultured under CTRL and TT04-treated conditions. (E, F) Representative immunofluorescence images and quantitative analysis of YAP and TFAP2C in E7.0 embryos cultured CTRL and TT04 treated conditions. (G, H) Representative immunofluorescence images and quantitative analysis of p-YAP and global O-GlcNAc in E7.0 embryos cultured CTRL and TT04-treated conditions. Data are presented as mean ± SEM. n≥3 independent biological replicates, with ≥5 blastocysts per group. Statistical significance was determined using a two-tailed unpaired Student’s t-test. ns, not significant; **P < 0.01; ****P < 0.0001. Scale bars, 100 μm.

Given the reduction in CDX2 expression, we next examined the Hippo signaling pathway. TT04-mediated O-GlcNAc inhibition significantly attenuated total YAP protein abundance without altering TFAP2C expression (Fig. 4E, F). Furthermore, TT04 treatment triggered a marked increase in phosphorylated YAP (p-YAP) levels (Fig. 4G, H), indicating enhanced YAP phosphorylation and cytoplasmic retention upon O-GlcNAc inhibition. Collectively, these findings demonstrate that O-GlcNAc acts as a critical metabolic signal that preserves TE identity by stabilizing the YAP protein.

### PPP–mTOR Signaling Axis Regulates TFAP2C

Glucose-derived nucleotides synthesized via the PPP serve as metabolic signals that activate mTOR signaling [5, 28]. To evaluate mTORC1 activity under glucose deprivation or PPP inhibition, we used phosphorylated ribosomal protein S6 (pS236-RPS6) as a downstream readout. IF analysis revealed a marked reduction in pS236-RPS6 levels in both G− and 6-AN–treated blastocysts compared to CTRL embryos (Fig. 5A–D). These results indicate that PPP-dependent metabolic flux is essential to sustain mTORC1 signaling during bovine blastocyst development.

**Fig 5.**
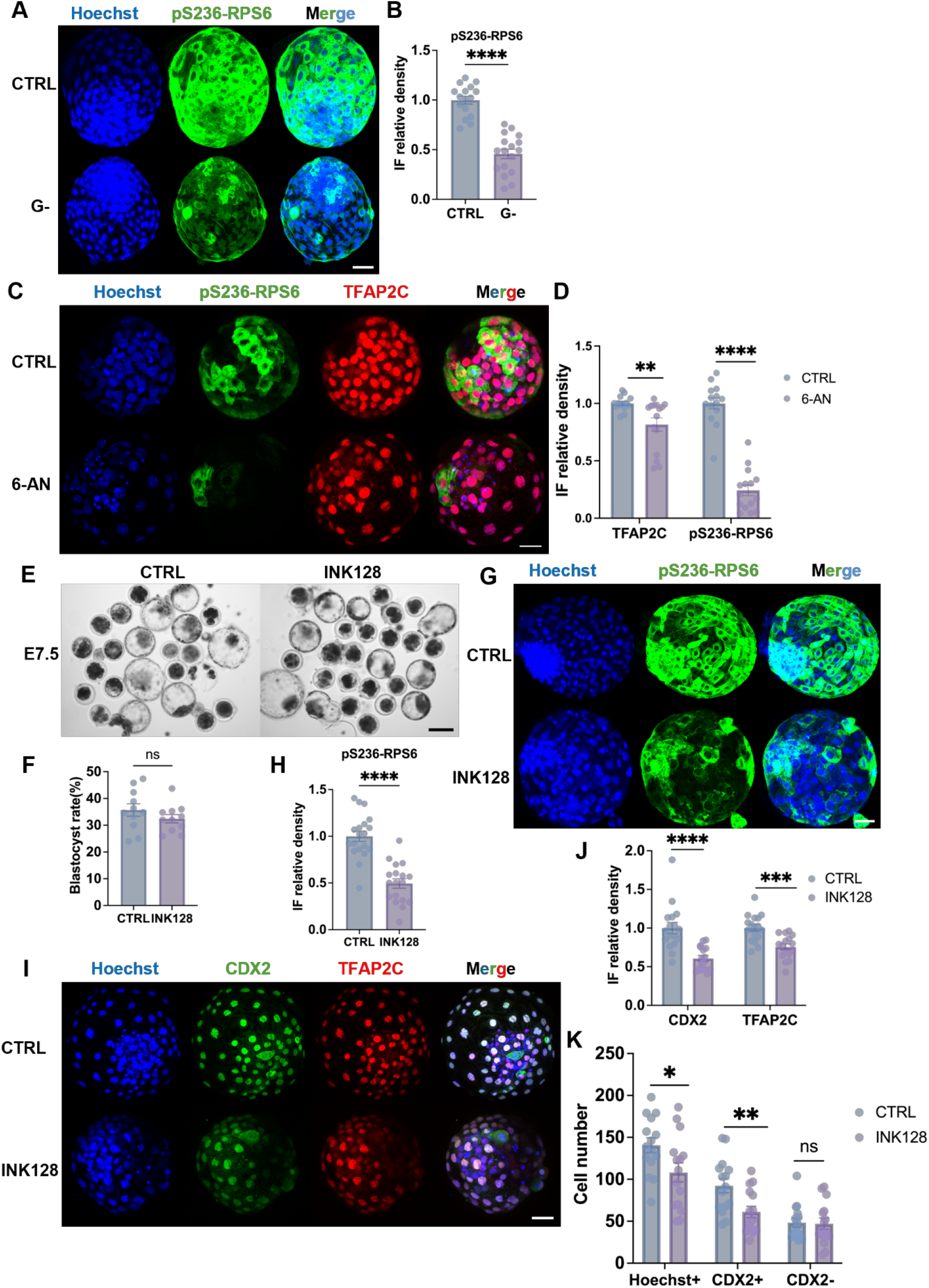
Inhibition of the PPP suppresses mTOR activity and disrupts TE differentiation. (A, B) Representative immunofluorescence images and quantitative analysis of pS236-RPS6 in E7.0 embryos cultured under CTRL or G– conditions. (C, D) Representative immunofluorescence images and quantitative analysis of pS236-RPS6 and TFAP2C in E7.5 embryos cultured under CTRL or 6-AN-treated conditions. (E, F) Representative bright-field images and quantification of blastocyst formation rates in E7.0 embryos cultured under CTRL or INK128-treated conditions. Scale bar, 50 μm. (G, H) Representative immunofluorescence images and quantitative analysis of pS236-RPS6 in E7.5 embryos cultured under CTRL or INK128-treated conditions. (I–K) Representative immunofluorescence images and quantitative analysis of CDX2 and TFAP2C in embryos under CTRL or INK128-treated conditions. Data are presented as mean ± SEM. n ≥3 independent biological replicates, with ≥5 embryos per groups. Statistical significance was determined using a two-tailed unpaired Student’s t-test: ns, not significant; *P < 0.05; **P < 0.01; ***P < 0.001; ****P < 0.0001. Scale bar, 100 μm

Given that mTOR promotes mRNA translation through the phosphorylation of RPS6 [5, 10, 11], we next interrogated whether the PPP regulates TFAP2C expression through an mTOR-dependent mechanism. Embryos were treated with the mTOR inhibitor INK128 from E0, which effectively suppressed mTORC1 activity, as evidenced by reduced pS236-RPS6 levels (Fig. 5G, H). While mTOR inhibition did not arrest overall blastocyst formation (Fig. 5E, F), it resulted in the significant downregulation of TFAP2C and CDX2 protein levels (Fig. 5I, J). Quantitative analysis further demonstrated a reduction in total blastocyst cell numbers following INK128 treatment, which was driven primarily by a selective loss of TE (CDX2^+^) cells, whereas the ICM (CDX2^−^) population remained unaffected (Fig. 5I, K). Collectively, these findings identify mTORC1 as a critical glucose-responsive signaling node that bridges PPP-derived nucleotide metabolism with the maintenance of TFAP2C expression and the expansion of the TE lineage.

### Single-cell transcriptomic profiling reveals lineage-specific metabolic signatures in bovine blastocysts

To further delineate the metabolic dynamics during bovine preimplantation development and corroborate our in vitro findings, we analyzed single-cell transcriptomic data spanning from the morula to the late blastocyst stages. UMAP dimensionality reduction successfully reconstructed the developmental trajectory and clearly segregated the distinct early embryonic cell lineages, including the TE, ICM, EPI, and PE (Fig. S4A). Heatmap analysis of metabolic gene modules revealed that oxidative phosphorylation (OXPHOS), glycolysis, the HBP, and the PPP exhibit distinct stage- and lineage-specific expression patterns during early embryogenesis (Fig. S4C and Fig 6 A, D, G). To robustly quantify these pathway-level changes and overcome the sparsity of single-cell data, we calculated metabolic module scores, which reflect the overall transcriptional enrichment of these entire metabolic networks within individual cells.

**Fig 6.**
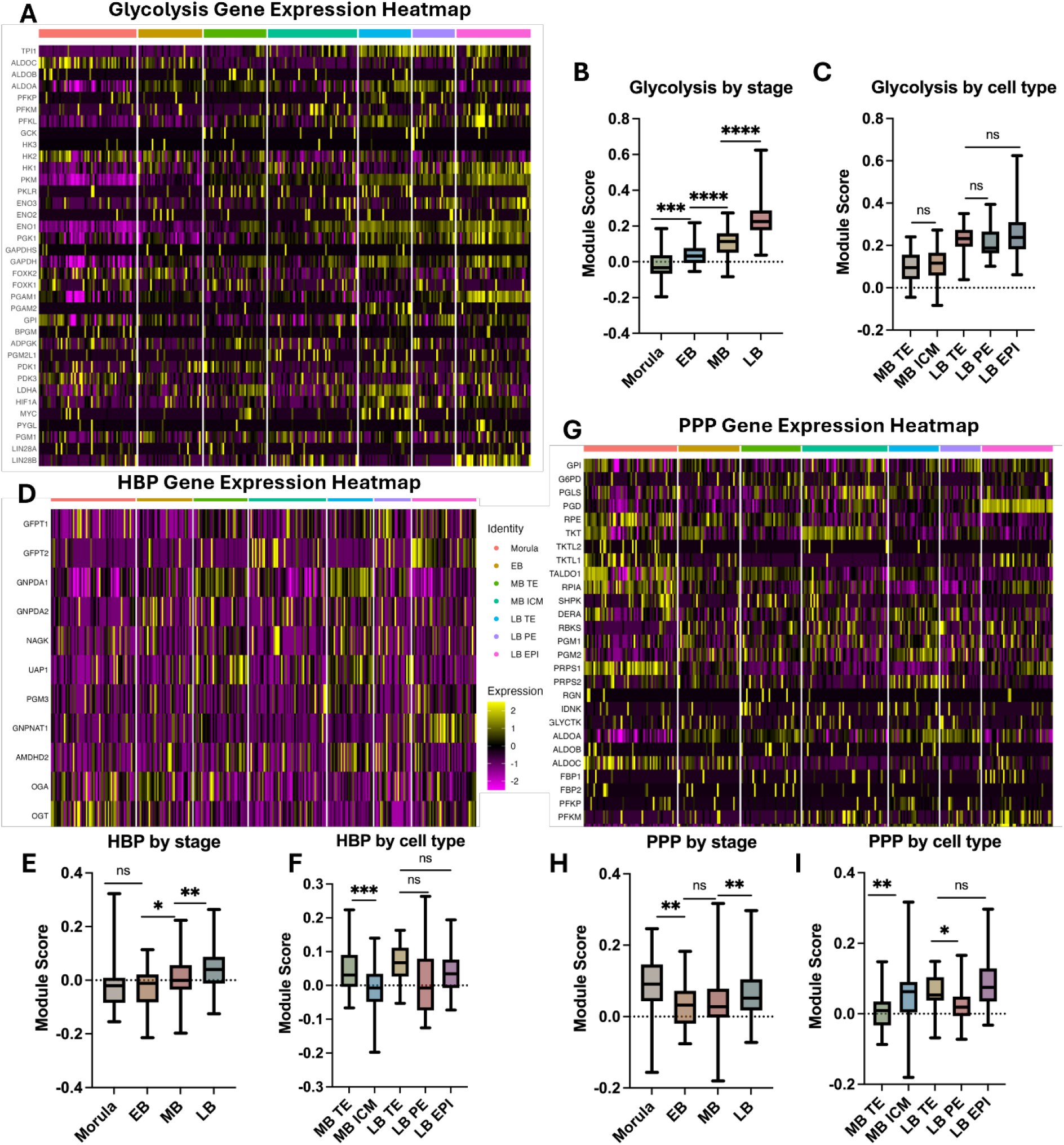
Quantitative analysis of metabolic pathway module scores during bovine preimplantation development. (A, D, G) Heatmap showing the expression of glycolysis,HBP and PPP module genes. (B, E, H) Bar graphs comparing the dynamic changes in module scores of glycolysis, HBP and PPP across grouped developmental stages. (C, F, I) Bar graphs illustrating the differences in module scores among TE, ICM, PE and EPI at the mid- and late-blastocyst stages. Data are presented as mean ± SEM. Statistical significance was determined using Welch’s one-way ANOVA, followed by Dunnett’s T3 post-hoc test: ns, not significant; *P < 0.05; **P < 0.01; ***P < 0.001; ****P < 0.0001.

To establish a macroscopic understanding of the metabolic landscape, we first evaluated the comprehensive module scores across all identified developmental stages and cell lineages (Fig. S4B). This overview revealed that OXPHOS maintains a consistently high and relatively stable transcriptional baseline throughout preimplantation development, likely fulfilling the basal energetic demands of the embryo. In contrast, the glucose-driven pathways displayed lower overall absolute scores but exhibited highly dynamic fluctuations, prompting us to further dissect their stage- and lineage-specific variations (Fig. S4B).

We next quantitatively assessed the dynamic shifts in these metabolic modules as embryos progressed through preimplantation development. Bar graphs comparing grouped stages revealed that during the transition from the morula to the mid-blastocyst (MB) stage, the expression modules for both OXPHOS and glycolysis generally increased (Fig. S4D and Fig 6B). However, at the late blastocyst (LB) stage, OXPHOS scores underwent a significant decline, whereas glycolysis continued to sharply rise (Fig. S4D and Fig 6B). This divergent pattern indicates a fundamental metabolic switch in energy production strategies, transitioning toward a predominantly glycolytic state, which is highly consistent with previously reported metabolic shifts in mammalian embryogenesis [21, 29-31].

In contrast to the continuous trajectory of glycolysis, the dynamic changes of the glucose-driven anabolic branches, HBP and PPP, were more complex. The HBP score initially exhibited a transient dip at the early blastocyst (EB) stage before steadily upregulating and peaking at the LB stage (Fig. 6F). Similarly, the PPP module showed a high baseline at the morula stage, followed by a decline at the EB stage, and a subsequent recovery and continuous increase through the MB and LB stages (Fig. 6H). Collectively, these results indicate that glucose-driven metabolic pathways undergo pronounced fluctuations during the morula-to-blastocyst transition.

Importantly, we investigated whether these glucose-driven pathways exhibited lineage-specific partitioning corresponding to our in vitro phenotypes. Quantitative comparison of module scores revealed highly nuanced, pathway-specific metabolic wiring among the cell lineages at the mid- and late-blastocyst stages (Fig. S4B). Notably, the glycolysis scores showed no significant differences between the TE and ICM lineages (Fig. 6C). This finding perfectly aligns with our earlier in vitro observation that inhibiting high-flux glycolysis with YZ9 does not selectively impair TE specification (Fig. S2K, M), confirming that basal glycolysis is broadly utilized rather than acting as a lineage-determining driver.

In stark contrast, the HBP module was robustly and specifically enriched in the TE compared to the ICM at the critical mid-blastocyst stage (Fig. 6F). This striking differential expression provides compelling in vivo transcriptomic evidence that HBP activity is preferentially partitioned to the TE, mechanistically explaining why TE specification relies heavily on the HBP-O-GlcNAc-YAP signaling axis.

In addition, the PPP module displayed high activity in both embryonic lineages. Notably, the module score in the MB ICM was significantly higher than that in MB TE, whereas LB TE scores were significantly higher than LB PE (Fig. 6I). This dynamic lineage-specific shift suggests that both ICM and TE rely on PPP during blastocyst development, albeit with different temporal requirements. The elevated PPP activity likely reflects the increased demand for nucleotide biosynthesis and redox homeostasis required to sustain rapid cellular proliferation [4, 5, 32, 33]. Consequently, inhibition of PPP is expected to affect both lineages and may therefore produce more severe developmental phenotypes than inhibition of other glucose-derived pathways. This robust and ubiquitous expression pattern perfectly aligns with our in vitro observations: pharmacological suppression of the PPP using 6-AN starting from the zygote stage results in a catastrophic developmental arrest at the morula stage (Fig S1H, I).

## Discussion

In this study, we demonstrate that bovine embryos exhibit a heightened tolerance to glucose deprivation compared to murine models. Although the absence of glucose significantly reduced blastocyst formation rates and compromised overall embryo quality, a subset of embryos retained the capacity to progress beyond the morula stage and successfully cavitate (Fig. 1A, B). This stands in stark contrast to mouse embryos, which undergo irreversible developmental arrest at the morula stage upon glucose withdrawal [2, 5]. In mice, this arrest is primarily attributed to a failure in the glucose-dependent Hippo-YAP axis to regulate CDX2, a factor essential for blastocoel formation in that species [2, 5]. However, while the Hippo pathway–dependent regulation of CDX2 is conserved in bovine embryos [23], TE differentiation is not exclusively dependent on the CDX2 regulatory circuit [14, 34]. Consequently, we observed that while glucose deficiency markedly attenuates the Hippo-YAP axis (Fig. 2G–L) and the expression of core TE markers CDX2 and GATA3 (Fig. 2A, B; Fig. S2A, B), bovine embryos remain morphologically capable of cavitation.

This phenotypic resilience is also likely underpinned by the superior metabolic flexibility of bovine embryos. Unlike murine embryos, which possess limited endogenous energy stores, bovine embryos are characterized by abundant lipid droplets that serve as alternative energy and biosynthetic substrates during nutrient stress [25, 35, 36]. Furthermore, the BO-SOF culture medium utilized in this study provides exogenous pyruvate, lactate, and amino acids, which likely act as a metabolic buffer, sustaining basal development even in the absence of glucose [37-39].

Similarly, human embryos exhibit metabolic plasticity and can progress to the blastocyst stage in glucose-deprived conditions [40, 41]. This divergence highlights species-specific metabolic coping strategies and underscores that the bovine model often recapitulates human embryonic development more faithfully than murine models. Therefore, our research model is designed to address the inherent translational limitations of the mouse model.

Our findings demonstrate that the three primary branches of glucose metabolism fulfill distinct functional roles during TE differentiation. We observed that basal OCR and exogenous pyruvate uptake remained stable in G− embryos (Fig. 3C–F), and inhibiting glycolytic flux with YZ9 failed to impair blastocyst formation or TE specification (Fig. 1E, F). These results indicate that glucose-derived pyruvate production is redundant for early lineage commitment in bovine embryos. This aligns with isotope tracing studies in mice showing that the carbon contribution of glucose to the pyruvate pool is minimal [5], as well as our metabolomics data showing stable intracellular pyruvate levels in G− blastocysts (Fig. S3A). Collectively, these findings reinforce the conclusion that the requirement for glucose in early embryogenesis is decoupled from bioenergetic production, serving instead to sustain critical biosynthetic flux and metabolic signaling [5, 8, 17].

Our results uncover a striking divergence in the requirement for the HBP between bovine and murine embryos. In mice, pharmacological inhibition of either the HBP (via DON) or its downstream effector, OGT (via TT04), triggers a terminal arrest at the morula stage [5]. In contrast, bovine embryos treated with TT04 successfully undergo cavitation despite a significant loss of O-GlcNAcylation (Fig. 3G, H). However, the terminal arrest observed following DON treatment (Fig. S1H, J) indicates that HBP-derived flux remains a fundamental requirement for bovine survival, likely providing substrates for N-glycan synthesis or other vital biosynthetic processes independent of O-GlcNAcylation.

Mechanistically, our data suggests a unique relationship between glycosylation and YAP stability. In murine embryos, O-GlcNAc is proposed to stabilize nuclear YAP independent of its phosphorylation status [5, 8, 42]. However, we observed that O-GlcNAc inhibition via TT04 significantly increased p-YAP levels (Fig. 4G, H), while reducing nuclear YAP accumulation and CDX2 expression (Fig. 4A–F). The relationship between glycosylation and YAP phosphorylation during embryogenesis is largely unexplored. However, mutating the Thr241 O-GlcNAc site significantly compromises YAP stability while promoting its phosphorylation in tumor cells [43]. We therefore speculate that inhibition of O-GlcNAc promotes YAP retention in the cytoplasm in bovine embryos, where it becomes increasingly phosphorylated.

Interestingly, unlike HBP-specific inhibition, glucose deprivation in bovine embryos resulted in a marked reduction in p-YAP (Fig. 1I, 1J). In cancer cells, AMPK-mediated site-specific phosphorylation serves as a primary mediator of glucose-deficiency signaling [44, 45]. Thus, we speculate that glucose deprivation reduces overall phosphorylation levels, leading to decreased p-YAP.

The role of glucose in regulating phosphorylation is further supported by its impact on the mTOR pathway [45]. We found that glucose deprivation significantly decreased phosphorylation of RPS6, a downstream target of mTORC1 (Figure 5A, B). In addition, inhibition of PPP disrupted mTOR signaling, and both PPP inhibition and mTOR suppression markedly reduced the expression of the TE-associated transcription factor TFAP2C (Fig. 5G-K). These findings indicate that PPP activity is essential for maintaining the mTORC1–TFAP2C signaling axis required for proper TE differentiation. Furthermore, inhibition of PPP from the zygote stage resulted in developmental arrest at the morula stage, which could not be rescued by supplementation with downstream metabolites (Fig. S1H–K). Glucose flux through the PPP serves a critical antioxidant function via NADPH production [32, 33]. When this pathway is disrupted at the G6PD level, murine embryos experience elevated reactive oxygen species (ROS) levels [5, 32]. While uridine can mitigate some defects, full phenotypic restoration requires a combinatorial approach addressing both nucleotide depletion and oxidative stress [5]. We thus speculate that the irreversible arrest observed in bovine embryos is due to elevated ROS levels induced by pathway inhibition, ultimately leading to embryonic lethality.

Beyond direct signaling cascades, glucose availability profoundly shapes the embryonic epigenetic landscape. We observed that disrupted glucose metabolism led to significant reductions in arginine methylation (ADMA and SDMA) alongside a pronounced decrease in the repressive histone modification H3K27me3 (Fig. 3I–N and Fig. S3E–H). Protein arginine methyltransferase 1 (PRMT1) accounts for approximately 85% of ADMA formation in cells [46, 47]. By inhibiting phosphorylation-dependent cytoplasmic retention and proteasomal degradation, PRMT1-mediated methylation promotes YAP stabilization and nuclear accumulation [48]. Therefore, we speculate that glucose may interfere with the Hippo pathway by affecting arginine methylation, thereby influencing TE differentiation. Collectively, our findings highlight a potential link between glucose metabolism and epigenetic control, providing a new avenue for investigating the mechanisms underlying lineage specification.

## Acknowledgements

We thank Prof. Henry J Leese for his helpful discussion.

## Author contributions

J.Q. conceived the project, designed and performed all experiments, analysed and interpreted the data, and drafted and critically revised the manuscript. R.S. secured funding for the project and critically revised the manuscript.

