## supplemental Figures and tables for "Glucose-dependent signalling pathways regulate TE differentiation in bovine embryos"

### Supplementary figures

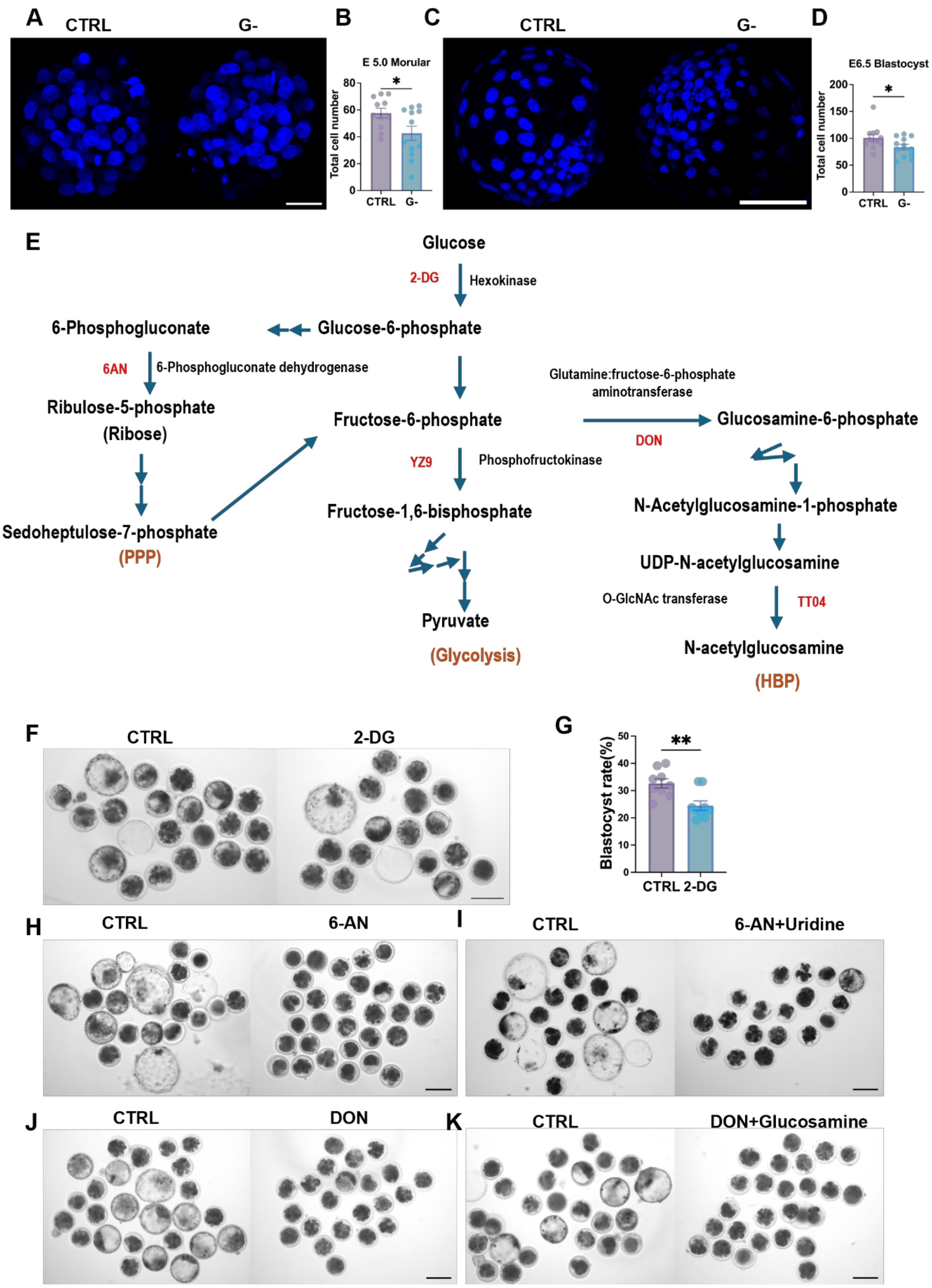


**Fig S 1 Impact of glucose deprivation and metabolic pathway inhibition on bovine embryo development.**

(A-D) Representative fluorescence images and quantification of total cell number in E5.0 morulae and E6.5 blastocyst cultured in CTRL and G- media. Nuclei are labeled with Hoechst33342 (blue). Three biological replicates, with at least three embryos per replicate.

(E) Schematic diagram illustrating key glucose metabolic pathways and the target sites of inhibitors.

(F, G) Representative bright-field images and corresponding blastocyst rate of embryos cultured with the glucose metabolism inhibitor 2-DG, relative to CTRL. Nine biological replicates, with at least 16 embryos per replicate.

(H, I) Representative images showing embryos treated with the PPP inhibitor 6-AN and the developmental rescue of 6-AN–treated embryos following uridine supplementation. At least three biological replicates, with at least 15 embryos per replicate.

(J, K) Representative images showing embryos treated with the HBP inhibitor DON and the developmental rescue of DON-treated embryos upon supplementation with Glucosamine. At least three biological replicates, with at least 15 embryos per replicate.

Data are presented as mean ± SEM. Statistical significance was determined using a two-tailed unpaired Student's t-test: ns, not significant; $*P < 0.05$; $**P < 0.01$. Scale bars: 100 μm (in A, C); 50 μm (in F, H, I, J, K).

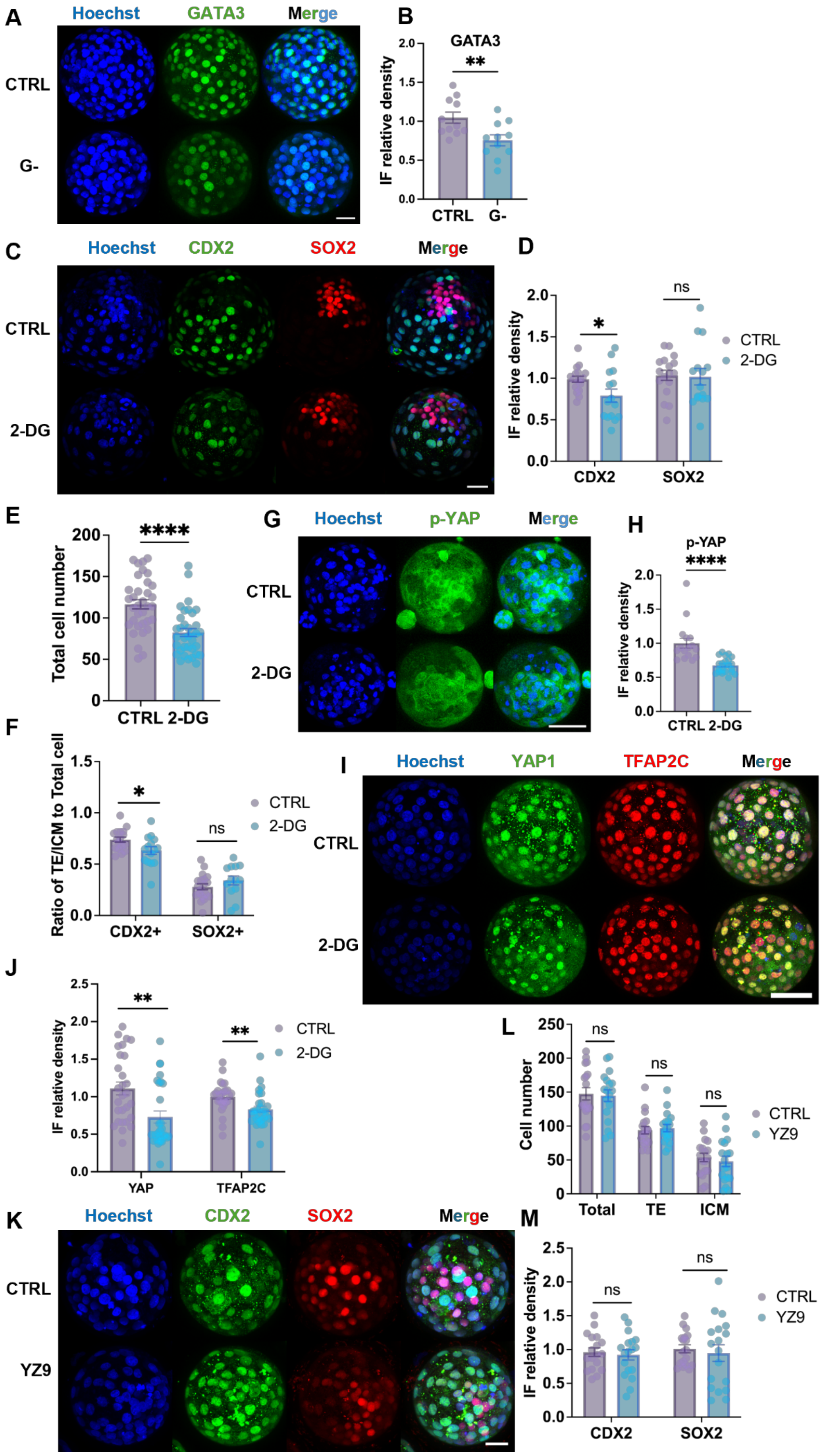
A
Hoechst
GATA3
Merge
CTRL
G-
B
GATA3
**
IF relative density
CTRL
G-
C
Hoechst
CDX2
SOX2
Merge
CTRL
2-DG
D
IF relative density
*
ns
CTRL
2-DG
CDX2
SOX2
E
Total cell number
****
CTRL 2-DG
G
Hoechst
p-YAP
Merge
CTRL
2-DG
H
p-YAP
****
IF relative density
CTRL 2-DG
F
Ratio of TE/ICM to Total cell
*
ns
CTRL
2-DG
CDX2+
SOX2+
I
Hoechst
YAP1
TFAP2C
Merge
CTRL
2-DG
J
IF relative density
**
**
CTRL
2-DG
YAP
TFAP2C
L
Cell number
ns
ns
ns
CTRL
YZ9
Total
TE
ICM
K
Hoechst
CDX2
SOX2
Merge
CTRL
YZ9
M
IF relative density
ns
ns
CTRL
YZ9
CDX2
SOX2

673

#### Fig S 2 Inhibition of glucose metabolism impairs TE specification through suppression of Hippo signalling in bovine embryos

(A, B) Immunofluorescence staining and quantification of GATA3 in E7.0 bovine blastocysts cultured CTRL or G– medium.

(C–F) Immunofluorescence staining and quantification of CDX2 and SOX2 in embryos cultured with or without 2-DG. Total cell number, ($CDX2^{+}$/$Hoechst^{+}$) and ICM ($SOX2^{+}$/$Hoechst^{+}$) cell ratios, and relative fluorescence intensities are shown.

(G, H) Immunofluorescence staining and quantification of p-YAP in CTRL and 2-DG–treated embryos.

(I, J) Immunofluorescence staining and quantification of YAP1 and TFAP2C in CTRL and 2-DG–treated embryos.

(K–M) Immunofluorescence staining and quantification of CDX2 and SOX2 in E7.0 embryos cultured in CTRL or YZ9 medium.

Data are presented as mean ± SEM from three biological replicates, with at least five blastocysts analyzed per replicate. Statistical significance was determined using a two-tailed unpaired Student's t-test: ns, not significant; $^{*}P < 0.05$; $^{**}P < 0.01$; $^{****}P < 0.0001$. Scale bars: 100 μm.

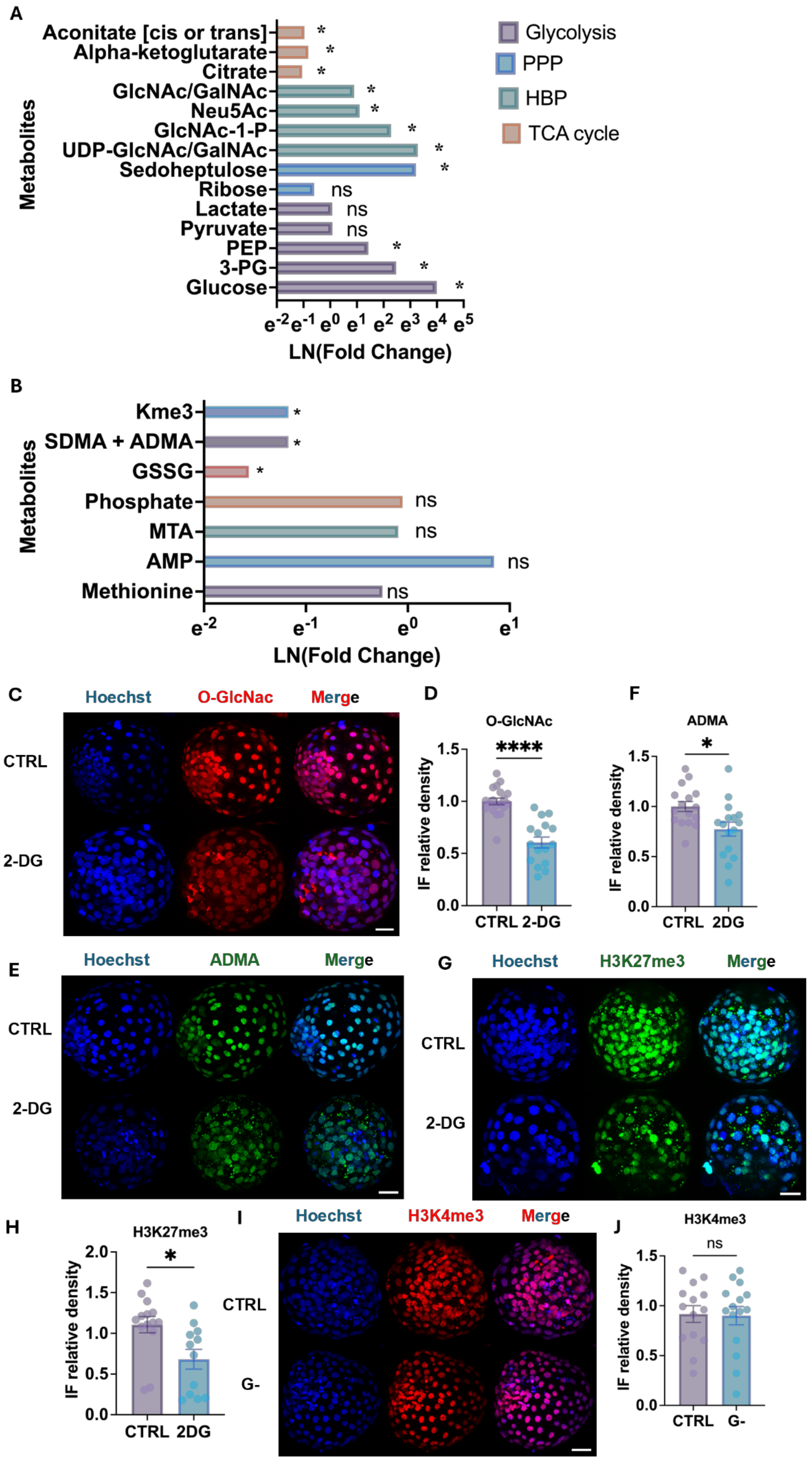
A
Aconitate [cis or trans]
Alpha-ketoglutarate
Citrate
GlcNAc/GalNAc
Neu5Ac
GlcNAc-1-P
UDP-GlcNAc/GalNAc
Sedoheptulose
Ribose
Lactate
Pyruvate
PEP
3-PG
Glucose
Metabolites
ns
*
e-2 e-1 e0 e1 e2 e3 e4 e5
LN(Fold Change)
Glycolysis
PPP
HBP
TCA cycle
B
Kme3
SDMA + ADMA
GSSG
Phosphate
MTA
AMP
Methionine
Metabolites
e-2 e-1 e0 e1
LN(Fold Change)
C
Hoechst
O-GlcNac
Merge
CTRL
2-DG
D
O-GlcNAc
****
IF relative density
1.5
1.0
0.5
0.0
CTRL 2-DG
F
ADMA
*
IF relative density
CTRL 2DG
E
Hoechst
ADMA
Merge
CTRL
2-DG
G
Hoechst
H3K27me3
Merge
CTRL
2-DG
H
H3K27me3
*
2.0
IF relative density
CTRL 2DG
I
Hoechst
H3K4me3
Merge
CTRL
G-
J
H3K4me3
ns
IF relative density
CTRL G-

**Fig S 3 Glucose deprivation reshapes carbon metabolism and epigenetic modifications in bovine blastocysts**

(A) Re-analysis of metabolomics data showing changes in carbon-associated metabolites following glucose perturbation. Metabolites involved in glycolysis, PPP, HBP, and TCA cycle are displayed. Data are presented as ln (fold change) relative to control conditions, highlighting pathway-specific alterations in glucose-derived carbon flux.

(B) Re-analysis of metabolomics data showing changes in selected amino acid– and methylation-related metabolites, including methionine cycle–associated intermediates, arginine methylation products (SDMA + ADMA), redox-related metabolites, and nucleotide-associated species. Data are presented as ln (fold change) relative to control conditions.

(C-J) Representative immunofluorescence images and corresponding quantification of O-GlcNAc, ADMA, H3K27me3, and H3K4me3 in blastocysts under G-, 2-DG or CTRL conditions. Nuclei were counterstained with Hoechst33342 (blue). Bar graphs show the relative immunofluorescence density. Scale bars: 100 μm.

Data are presented as mean ± SEM from three biological replicates, with at least five blastocysts analyzed per group. Statistical significance was determined using a two-tailed unpaired Student's t-test: ns, not significant; $^*P < 0.05$; $^{****}P < 0.0001$. Scale bars: 100 μm.

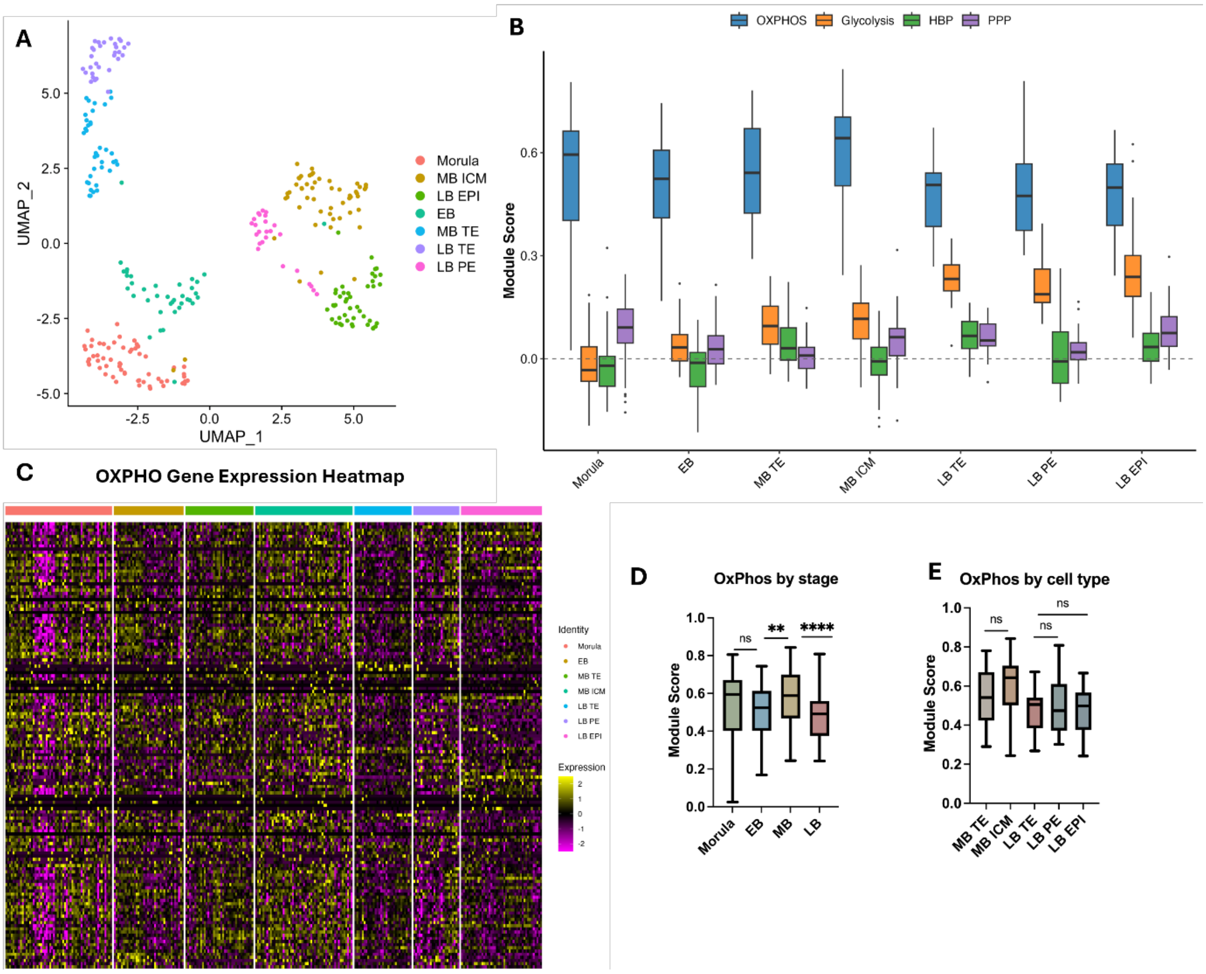


**Fig S4 Single-cell profiling of cell lineages and metabolic states in early bovine embryogenesis**

(A) UMAP dimensionality reduction and clustering analysis of bovine preimplantation embryonic cell lineages, showing the developmental trajectory from the morula to the late blastocyst stage. Abbreviations: EB, early blastocyst; MB, mid blastocyst; LB, late blastocyst; TE, trophectoderm; ICM, inner cell mass; PE, primitive endoderm; EPI, epiblast.

(B) Boxplots showing the comprehensive module scores for OXPHOS, Glycolysis, HBP, and PPP across different developmental stages and defined cell lineages.

(C) Heatmap showing the expression of OXPHOS module genes.

(D) Bar graphs comparing the dynamic changes in module scores of OXPHOS across grouped developmental stages.

(E) Bar graphs illustrating the differences in module scores among TE, ICM, PE and EPI at the mid- and late-blastocyst stages.

Data are presented as mean ± SEM. Statistical significance was determined using Welch's one-way ANOVA, followed by Dunnett's T3 post-hoc test: ns, not significant; $^{**}P < 0.01$; $^{****}P < 0.0001$.

**Table S1 Chemical information**

| | Concentration | Supplier | CatLog NO. |
|---|---|---|---|
| 2-Deoxy-D-glucose (2-DG) | 20 μM | Sigma-Aldrich | D8375 |
| 6-Diazo-5-oxo-L-norleucine (DON) | 10 μM | Sigma-Aldrich | D2141 |
| 6-Aminonicotinamide (6-AN) | 2 μM | Sigma-Aldrich | A68203 |
| ST045849 (TT04) | 20 μM | Sigma-Aldrich | SML2702 |
| YZ9 | 10 μM | Sigma-Aldrich | SML1270 |
| INK128 | 100 nM | Cambridge Bioscience | CAY11811 |
| Uridine | 200/400 μM | Sigma-Aldrich | G4875 |
| D-(+)-Glucosamine hydrochloride | 250/500 μM | Sigma-Aldrich | U3003 |

**Table S2 Primary antibody information**

| Primary antibody | Concentration | Host | Supplier | CatLog NO. |
|---|---|---|---|---|
| Anti-NANOG | 1:100 | Mouse | Thermo Fisher | 14-5768-82 |
| Anti-GATA3 | 1:250 | Rabbit | Abcam | ab199428 |
| Anti-GATA6 | 1:1600 | Rabbit | Cell Signalling Technology | 5851T |
| Anti-SOX2 | 1:100 | Rat | Thermo Fisher | 14-9811-82 |
| Anti-ADMA | 1:200 | Rabbit | Cell Signalling Technology | 13522 |
| Anti-SDMA | 1:100 | Rabbit | Cell Signalling Technology | 13222 |
| Anti-CDX2 | 1:100 | Rabbit | Abcam | ab227201 |

| | | | | |
|---|---|---|---|---|
| Anti-H3K27me3 | 1:200 | Rabbit | Cell Signalling Technology | 9733 |
| Anti-H3K4me3 | 1:200 | Rabbit | Cell Signalling Technology | 9751T |
| Anti-O-GlcNAc | 1:100 | Mouse | Abcam | ab2739 |
| Anti-YAP | 1:100 | Rabbit | Cell Signalling Technology | 14074S |
| Anti-pYAP | 1:100 | Rabbit | Thermo Fisher | PA517481 |
| Anti-TFAP2C | 1:100 | Mouse | Santa Cruz | sc-12762 |
| Anti-TEAD4 | 1:100 | Mouse | Abcam | ab58310 |
| Anti-RPS6 | 1:100 | Rabbit | Cell Signalling Technology | 4858T |
| Anti- P4E-BP1 | 1:200 | Rabbit | Cell Signalling Technology | 2855T |

**Table S3 Secondary antibody information**

| Secondary antibody | Concentration | Host - Target | Supplier | CatLog # |
|---|---|---|---|---|
| Alexa Fluor™ 488 | 1:100 | Goat anti-Mouse | Thermo Fisher | A-11001 |
| Alexa Fluor™ 594 | 1:100 | Goat anti-Rat | Thermo Fisher | A-11007 |
| Alexa Fluor™ 488 | 1:100 | Goat anti-Rabbit | Thermo Fisher | A-11008 |
| Alexa Fluor™ 594 | 1:100 | Goat anti-Rabbit | Thermo Fisher | A-11012 |

| | | | | |
|---|---|---|---|---|
| Alexa Fluor™ 568 | 1:100 | Goat anti-Mouse | Thermo Fisher | A-11004 |
